# Genome structure and molecular phylogeny of the only Eurasian *Boechera* species, *Boechera falcata* (Brassicaceae)

**DOI:** 10.1101/2025.03.11.642644

**Authors:** Danil Zilov, Terezie Mandáková, Marina Popova, Michael D. Windham, Vladimir Brukhin

## Abstract

*Boechera falcata* (Turcz.) Al-Shehbaz (earlier known as *Arabis turczaninowii* Ledeb.) is a herbaceous perennial belonging to the East Siberian, boreal-steppe ecotype. It is the only species of the diverse genus *Boechera* that grows on the Eurasian continent, all others being endemic to North America and Greenland. In all likelihood, *Boechera falcata* or its ancestral lineage migrated from North America to Eastern Siberia across the Bering Land Bridge when it was exposed during the height of Pleistocene glaciation. *Boechera* is interesting in that many species in this genus are allodiploid and triploid apomicts that arose through complex hybridization of sexual species and ecotypes. To date, the genomes of only two American *Boechera* species, *B. stricta* and *B. retrofracta*, have been sequenced, analyzed, and published. In the present study, we sequenced, assembled to the chromosome level, and analyzed the *B. falcata* genome, which was highly homozygous with a size of 189.36 Mb. Molecular phylogenetic analysis of the nuclear and organelle genomes revealed a high degree of relatedness of *B. falcata* to North American relatives. Investigation of the structure of pachytene, mitotic, and diakinesis chromosome spreads (n = 14) using cytogenetic analysis and comparative chromosome painting (CCP) allowed us to identify all 22 genomic blocks of crucifers and found that five of the seven *B. falcata* chromosomes were collinear with their corresponding chromosomes in the ancestral Boechereae genome, and two chromosomes had undergone pericentric and paracentric inversions. Allelic analysis of the apomixis marker *APOLLO* gene revealed that *B. falcata* contains only sex alleles of this gene. The availability of the genome of the only Asian *Boechera* species will facilitate studies of the evolution and phylogeny of Brassicaceae species and apomixis.

## Introduction

As currently defined (Alexander et al. 2013; Hay et al. 2023), the genus *Boechera* Á. Löve & D. Löve (Brassicaceae) comprises c. 75 sexual diploid taxa and c. 355 genetically distinct hybrid lineages with distributions concentrated in western North America. *Boechera falcata* (Turcz.) (*=Arabis turczaninowii* Ledeb.), the sole species found in Eurasia, is endemic to Eastern Siberia and the Far East. Molecular marker data suggest its origin may be linked to the migration of ancestral forms from North America to Siberia over the Bering Land Bridge (Al-Shehbaz, 2005; Windham and Al-Shehbaz, 2007; Kiefer et al., 2009; Alexander et al., 2013; Brukhin et al., 2019). Although earlier classifications placed members of *Boechera* within the genus *Arabis* L. based on superficially similar morphological traits, karyological studies revealed chromosomal differences, prompting Á. Löve and D. Löve (1975) to separate the genus *Arabis* (s.l.) into New World species with x = 7 (now classified as *Boechera*) and Old World species with x = 8 (retained in *Arabis*). Subsequent molecular phylogenetic analyses have confirmed that these two genera belong to distinct clades within the Brassicaceae (Koch et al., 2000; Beilstein et al., 2010; Nikolov et al., 2019; Hendriks et al., 2023), demonstrating that their similarities are convergent rather than indicative of a close evolutionary relationship.

Species of the genus *Boechera* are closely related to the well-studied model plant *Arabidopsis* of the Brassicaceae family. *Boechera* is the only genus within the Brassicaceae family where apomixis is well documented (Böcher, 1951, 1969; Brukhin et al., 2019). Apomixis is asexual reproduction of plants through seeds, in which the embryo arises not from the fusion of male and female germ cells, as occurs in sexual reproduction, but is a genetic clone of the maternal plant and develops parthenogenetically. The ability to produce clones of a maternal plant and hence fix the desired genotypes in subsequent generations of various agricultural species will facilitate the development of effective plant breeding strategies based on the knowledge of the genetic aspects of apomixis, and understanding the molecular mechanisms underlying it has a great potential for application in agriculture due to its ability to fix valuable genetic traits of the maternal plant in an indefinite series of generations (Grossniklaus et al., 2002; Bicknell and Koltunow, 2004; Van Dijk, 2009; Brukhin, 2017). Allodiploid and triploid apomictic accessions of *Boechera* show signs of hybridogenic origin, due to which changes in their chromosome structure are observed with characteristic occurrence of alloploidy, aneuploidy, substitution of homologous chromosomes and the appearance of aberrant extra chromosomes (Mandáková et al., 2015; Brukhin et al., 2019). Therefore, species from the genus *Boechera* are unique models for molecular genetic studies of apomixis and the influence of hybridization of sexual forms on its occurrence.

To date, whole genome assemblies have been published for only two *Boechera* species: *B. stricta* (Lee et al., 2017) and *B. retrofracta* (Kliver et al., 2018). These species are of particular interest because, while they reproduce predominantly sexually, many of their hybrids are apomicts. The Far Eastern *Boechera falcata* studied here is a key outgroup species for understanding the genome structure of the genus and, possibly, the evolution of genetic loci associated with apomixis. This study aims to: 1) Achieve a chromosome-scale genome assembly of *B. falcata*. 2) Perform a phylogenetic analysis to explore its relationship with North American *Boechera* species. 3) Analyze genome structure, perform Hi-C genomic analysis technique, comparative genome analysis, phylogenetic analysis based on nuclear, and chloroplast DNA. 4) Study of the chromosome structure exploiting CCP (comparative chromosome painting) analysis and the location of ACK (Ancestral Crucifer Karyotype) within the chromosomes (as was described in Mandáková et al., 2015). 5) Study alleles of the apomixis-associated *APOLLO* locus (encoding NEN exonuclease) in *B. falcata* and add to the phylogenetic tree of *APOLLO* locus isoforms of the Boechera species published earlier (Kliver et al., 2018).

## Material and Methods

### 1. Plant Material

Seeds of *B. falcata* were collected from plants (Figure 1) in natural habitats: a petrophytic open community of *Artemisia gmelinii* Weber ex Stechm. and *B. falcata* on the southern slope with a steepness of 45° (Tenkinsky district of the Magadan region, 10 km southeast of the village of Orotuk - 62° 01’50.586” N, 148° 38’46.591” E).

**Figure 1.**
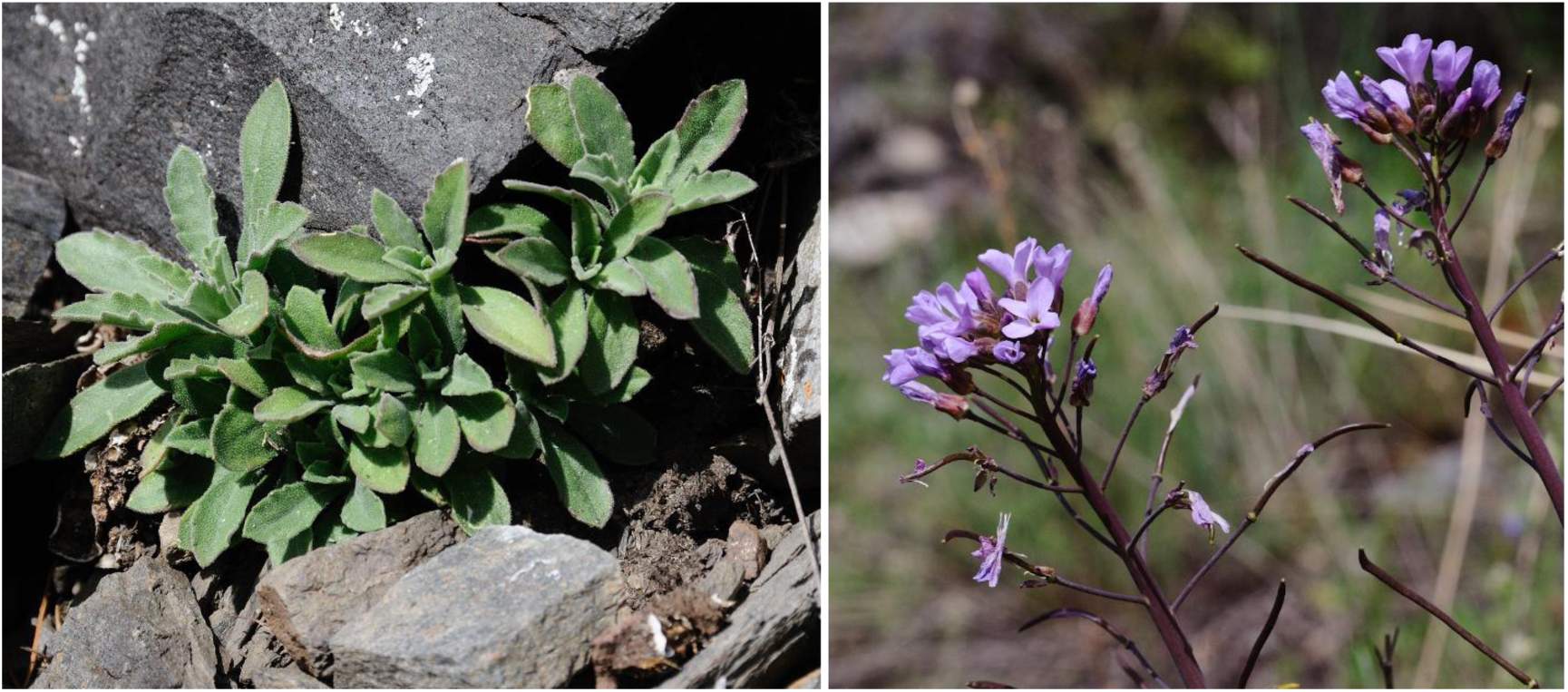
Plants of *Boechera falcata* (leave rosette and blooming plants with siliques) growing in the Kolyma River valley in the Magadan region, from which seeds were collected.

Plants were cultivated from seeds in the greenhouse of CEITEC Masaryk University under the standard conditions (16/8 h light/dark, 21/18 °C, 15 µmol/m2/s). Specimens are deposited at the Herbarium of Masaryk University (BRNU): BRNU sheet numbers: 696364 (https://brnu.jacq.org/BRNU696264) and 696267 (https://brnu.jacq.org/BRNU696267) (Suppl. Figure 1).

**Suppl. Figure 1.**
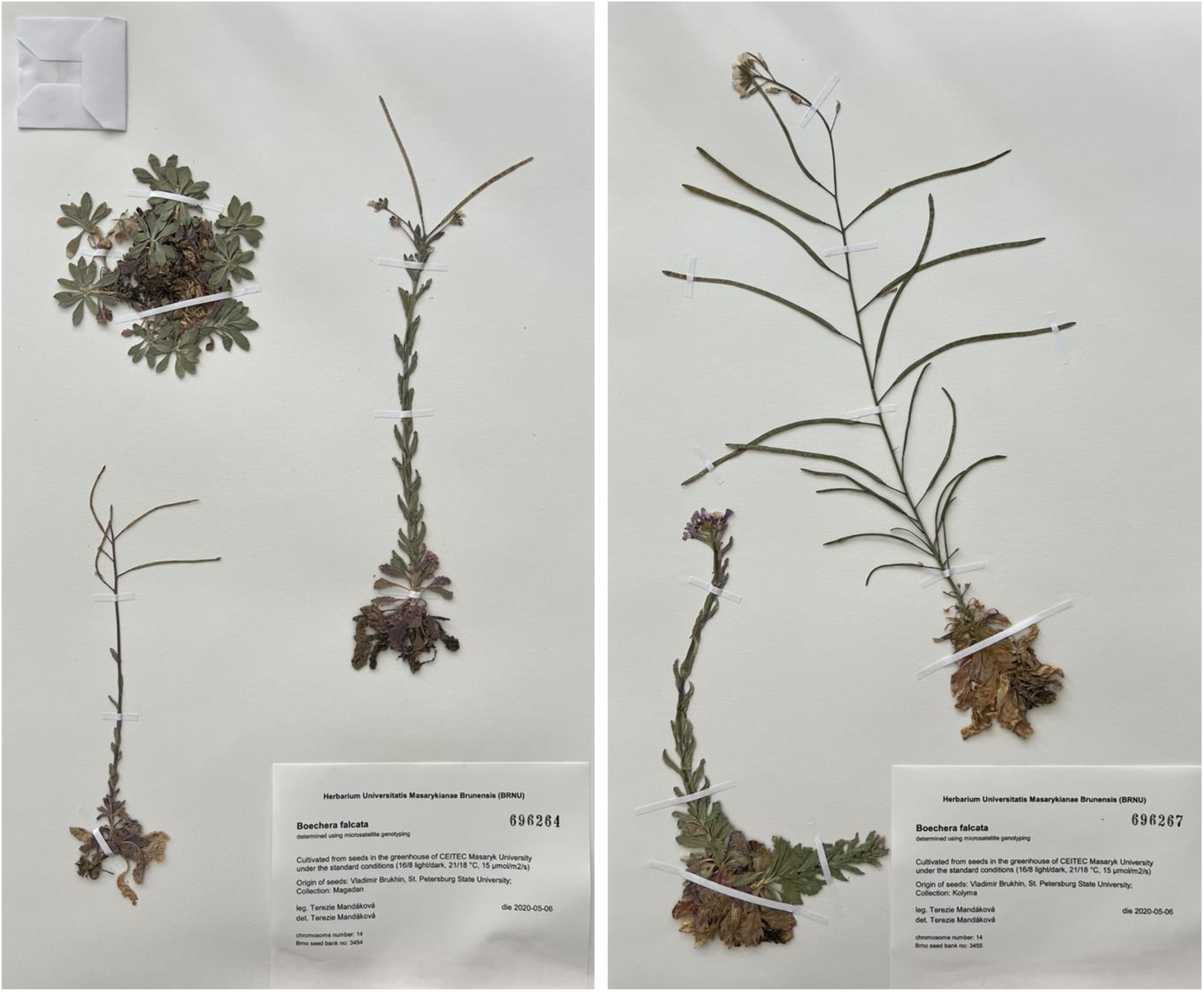
Herbarium specimen of *B. falcata* investigated, deposited in the herbarium of Masaryk University (BRNU).

### 2. Microsatellite genotyping

Genomic DNA was extracted using the Qiagen DNeasy Plant Mini Kit. Microsatellite allele variation was assessed at 15 previously published loci (A1, A3, B6, Bdru266, BF3, BF9, BF11, BF15, BF18, BF19, BF20, C8, E9, ICE3, ICE14) following Beck et al. (2012). Forward primers for each locus were labeled with either 6-FAM or HEX. Multiplex polymerase chain reaction (PCR) was performed for sets of three loci simultaneously. Each 50 μl reaction contained 15 μl of 2× Qiagen Multiplex PCR Master Mix, 3 μM of the primer set, and approximately 40 ng of gDNA template. The PCR conditions included an initial denaturation at 95°C for 15 min, followed by 30 cycles of denaturation at 95°C for 30 s, annealing at 53°C for 90 s, and extension at 72°C for 60 s, with a final extension step at 60°C for 30 min. PCR products were first analyzed on a 1–3% agarose gel. Capillary electrophoresis was then performed by Macrogen using a 400HD standard. Electropherograms were analyzed with the Microsatellite plugin of Geneious software, and species identity was determined using the Boechera Microsatellite Website https://windhamlab.biology.duke.edu/ (Li et al., 2017).

### 3. Chromosome preparation and comparative chromosome painting

Mitotic and meiotic (pachytene) chromosome preparations were performed according to the protocol outlined by Mandáková and Lysak (2016). Before analysis, suitable slides were pre-treated with RNase (100 mg/ml) and pepsin (0.1 mg/ml). Comparative chromosome painting was performed according to Mandáková et al. (2020). Fluorescent signals were examined and captured using a Zeiss Axioimager epifluorescence microscope equipped with a CoolCube camera (MetaSystems). Images were acquired separately for each of the four fluorochromes using their respective excitation and emission filters (AHF Analysentechnik). The collected images were then pseudocolored and merged using Adobe Photoshop CS6 (Adobe Systems).

### 4. Whole Genome Sequencing Strategy DNA extraction and prob preparation

DNA was extracted following Huang et al. (2023) protocol. The *B. falcata* genome was sequenced on the three WGS platforms: Illumina NovaSeq with 150bp PE sequencing run, final 75Gb data obtained, with 243x coverage; Pacbio sequel II (HiFi/CCS mode/cell) (1 SMRT cell), genome coverage - 129x; Hi-C genomic analysis technique ∼50M Read-Pairs, genome coverage - 30x.

#### Raw Data Filtration and Pre-Processing

Raw Illumina reads and HiC reads trimming were processed using Trimmomatic v0.32 (Bolger et al., 2014) with the following parameters ILLUMINACLIP:2:30:10:2:True LEADING:3 TRAILING:3 MINLEN:36. HiFi reads evaluation of length and quality stats was performed with pauvre tool (https://github.com/conchoecia/pauvre).

The quality assessment of HiFi reads using the Pauvre tool indicated a mean read length of 13,387 bases and an N50 value of 13,975 bases. The Phred quality scores ranged from 20 to 50, with a peak at 40 (Suppl. Figure 2).

**Suppl. Figure 2.**
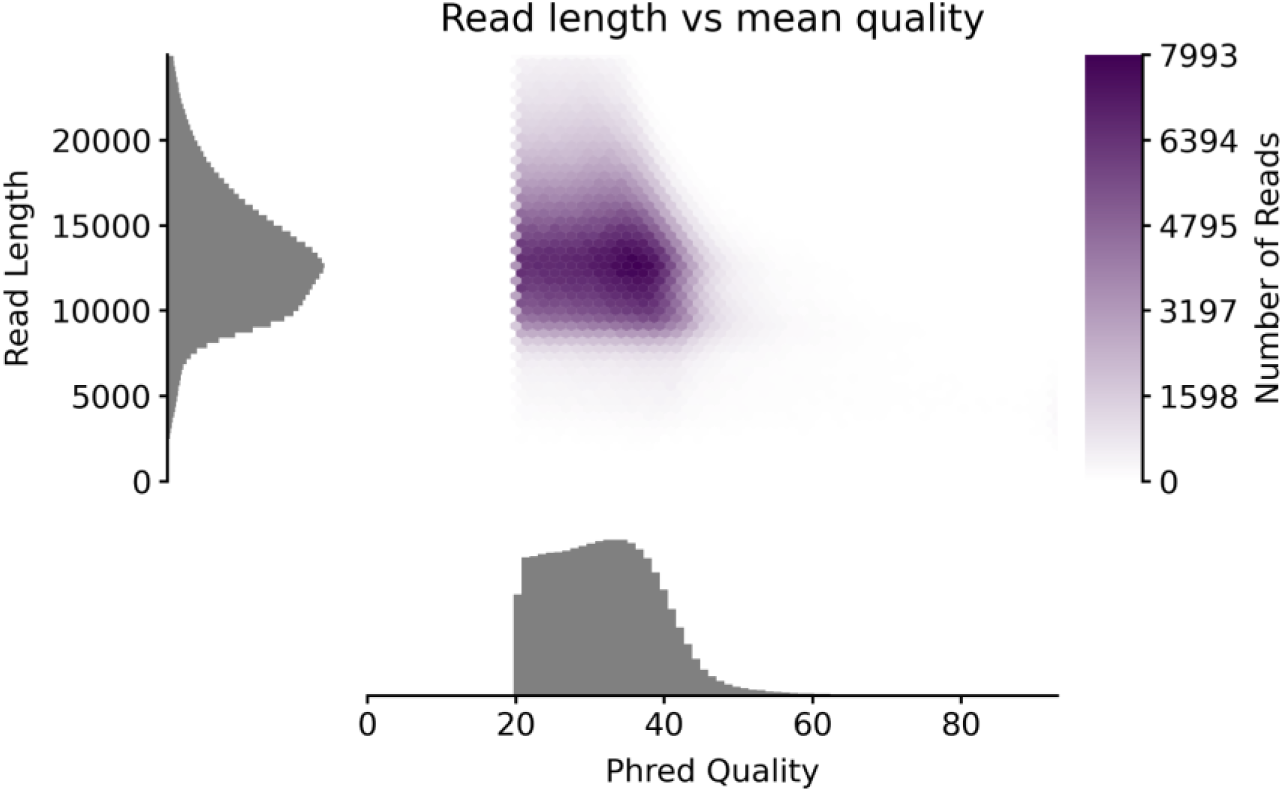
Distribution of read length and phred quality scores for the HiFi library.

#### Genome Coverage Estimation

Estimation of the genome coverage based on the 23-mer distribution and *k*-mer based statistics was performed using the a genomic k-mer counter Meryl (https://github.com/marbl/meryl), k-mer databases of HiFi and Illumina WGS reads, and GenomeScope2 (Ranallo-Benavidez et al., 2020).

### 5. Genome Assembly and Quality Control

For genome assembly we performed a bake off with four different assemblers: for HiFi reads: flye v2.9.2-b1786 (Kolmogorov et al., 2019), verkko v1.4.1 (Rautiainen et al., 2023), nextdenovo 2.5.2 (Hu et al., 2023), and hifiasm v0.19.6-r595 (Cheng et al., 2021) (Table 1). Flye was run with the --keep-haplotypes option, while verkko and nextdenovo targeted a genome size of 200Mbp. Nextdenovo and hifiasm were executed with standard parameters for HiFi reads. The assembly process was automated using the hifiblender pipeline [https://github.com/zilov/hifiblender].

**Table 1.** Comparative statistics of genome assembly by different assemblers.

|  | **NextDenovo** | **flye** | **hifiasm** | **verkko** |
| --- | --- | --- | --- | --- |
| Contigs № | 50 | 145 | 1232 | 2041 |
| Total length (bp) | 189363209 | 201258247 | 249420951 | 246125585 |
| GC (%) | 35.95 | 35.96 | 36.53 | 37.85 |
| N50 | 12407843 | 6341302 | 30472172 | 7370569 |
| L50 | 7 | 11 | 4 | 11 |
| K-mers not found in reads (missing) | 1929 | 2859 | 21778 | 75086 |
| K-mers overly represented in assembly | 1460638 | 3908427 | 31538183 | 27968096 |
| Merquy QV | 62.83 | 62.09 | 46.84 | 47.36 |
| Merfin QV* | 32.99 | 30.69 | 24.05 | 23.71 |
| Total low-coverage regions: | 0 | 0 | 0 | 0 |
| BUSCO | C:98.8%[S:95.8%, D:3.0%],F:0.2%,M:1.0%,n:4596 | C:98.8%[S:95.7%, D:3.1%],F:0.2%,M:1.0%,n:4596 | C:98.8%[S:95.9%, D:2.9%],F:0.2%,M:1.0%,n:4596 | C:98.7%[S:95.6%, D:3.1%],F:0.2%,M:1.1%,n:4596 |

Quality of the assembly was assessed using multiple metrics: N50, L50, BUSCO completeness, and QV (Quality Value). QUAST (Gurevich et al., 2013) was utilized for contiguity statistics, BUSCO (Seppey et al., 2019) for completeness assessment, and Merfin (Formenti et al., 2022) for QV (Quality Value) calculation using a Meryl HiFi + Illumina hybrid k-mer database (k=23). Coverage of the genome was calculated with GenomeScope2 (Ranallo-Benavidez et al., 2020).

Haplotype phasing was performed using hapdup [https://github.com/KolmogorovLab/hapdup] with default parameters. Polishing was conducted with nextpolish (Hu et al., 2020) using Illumina reads, with the number of steps (three for the first pseudo-haplotype, two for the second) determined by QV value progression checked on hybrid HiFi and Illumina Meryl database.

Scaffolding and genome curation Hi-C was facilitated by yahs (Zhou et al., 2023), followed by genome curation using juicebox (Robinson et al., 2018). The hic-scaffolder pipeline [https://github.com/zilov/hic-scaffolder] streamlined Hi-C read alignment and scaffolding processes. Finally, we aligned the orientation and order of chromosomes to match the *Boechera stricta* assembly (NCBI GenBank assembly ID: GCA_018361395.1).

### 6. Organelle DNA Assembly and Annotation

Mitochondrial and chloroplast genomes were assembled using contigs from the whole genome assembly after polishing and HiC scaffolding steps. For the mitochondrial genome, we employed MitoHiFi (Uliano-Silva et al., 2023), using the *B. stricta* mitochondrion as a reference to identify and assemble a complete circular mitochondrial sequence.

The chloroplast genome assembly was more complex. MitoHiFi was unable to resolve a circular molecule, so we used a manual approach. The *B. stricta* chloroplast genome served as a reference for identifying relevant contigs. We then performed a BLAST search to find intersections between these contigs. The chloroplast genome assembly required additional manual steps. While MitoHiFi software typically assembles circular organelle genomes, it failed to automatically reconstruct the complete circular chloroplast genome in our case. Therefore, we developed an alternative approach. First, we used the *B. stricta* chloroplast genome as a reference to identify chloroplast-derived contigs. Then, we performed BLAST analysis to find overlapping regions between these contigs. Using these overlapping sequences, we manually constructed the complete circular chloroplast genome.

Chloroplast and mitochondrial genome annotation and visualization were performed using GeSeq (Tillich et al., 2017) and manually curated by OGDRAW (Greiner et al., 2019), respectively.

### 7. Nuclear Genome Annotation

#### De novo Repeats Modeling With Repeatmodeler

To call for repetitive elements in the genome we used EarlGrey (Baril et al., 2024) repeat masking pipeline. A custom repeat database was created by combining the PlantRep database (Luo et al., 2022) and the nrTEplants dataset (Contreras-Moreira et al., 2021). Initial masking was conducted with the “eukarya” search term. De novo repeat identification performed with RepeatModeler (Flynn et al., 2020). The custom database was then used for subsequent rounds of masking to ensure identification of repetitive elements specific to plant genomes. EarlGrey utilizes a combination of tools, including RepeatModeler (Flynn et al., 2020), RepeatMasker [https://repeatmasker.org/], LTR_FINDER (Xu and Wang, 2007), and others to refine and cluster repeat sequences.

For coding gene annotation, we employed BRAKER v3.0.8 (Gabriel et al., 2024) in compleasm (Huang and Li, 2023) mode, utilizing the Viridiplantae OrthoDB (Kriventseva et al., 2019) protein database for homology-based prediction.

### 8. Comparative analysis of Boechera falcata and Boechera stricta genomes

We compared the assembled genome of *B. falcata* with the published *Boechera stricta* genome (NCBI GenBank assembly ID: GCA_018361395.1). Both genomes were evaluated using QUAST and BUSCO. QV calculations were performed differently: for *B. falcata*, k-mer analysis was carried out based on the hybrid HiFi and Illumina reads (https://github.com/arangrhie/T2T-Polish/tree/master/merqury), while for *B. stricta* k-mer analysis performed using only Nanopore reads due to their availability. Repeat masking and annotation were performed on both genomes. Synteny analysis between *B. falcata* and *B. stricta* chromosomes was conducted using SyRI (Goel et al., 2019). Whole-genome alignment visualization was generated with the help of D-GENIES (Cabanettes and Klop, 2018) to produce a comparative dotplot.

### 9. Phylogenetic Analysis of B. falcata Among the Other Boechera and Brassicaceae Species

The phylogenetic analysis of *B. falcata* was conducted in the context of other *Boechera* and Brassicaceae species using a combination of Angiosperms353 and Brassicaceae764 probe sets, following a methodology similar to that described by Hay et al. (2023). Loci were assembled using HybPiper version 2.1.7 with Illumina reads. Assembled loci were added to the existing locus files of other *Boechera* species. Each locus was aligned using MAFFT (Katoh and Toh, 2008) with the L-INS-I method. Alignments were concatenated into a supermatrix using FASconCAT-G (Kück and Longo, 2014) and subsequently trimmed using trimAl (Capella-Gutiérrez et al., 2009) with the -automated1 setting to reduce gaps and missing data. Phylogenetic tree construction was performed using RAxML-NG v1.2.0 (Kozlov et al., 2019), employing the GTR+F+R model and generating 1000 bootstrap replicates.

To analyze the phylogeny based on the chloroplast genome, we extracted 47 chloroplasts genes (listed in Suppl. Table 3) from the complete chloroplast assemblies of *Boechera* species and other Brassicaceae species reported elsewhere (Zhu et al., 2021). Phylogenetic analysis of chloroplast genomes was performed in MEGA (v.11.0.10): ClustalW(Codons) option with default settings was used for alignment of gene sequences. Phylogenetic tree was constructed with the Neighbor-Joining test with bootstrap 1000.

### 10. Analysis of APOLLO Sex- and Apo-alleles in B. falcata Genome

The *APOLLO* gene (Corral et al. 2013) was identified in the annotated genome using BLAST search. Additional gene sequences for phylogenetic analysis were obtained from Bakin et al. (2022). Multiple sequence alignment was performed using MUSCLE (Edgar et al., 2004). Phylogenetic tree construction was carried out with RAxML-NG v1.2.0 (Kozlov et al., 2019), employing the following parameters: -all -model JTT+G --bs-trees 1000 --tree pars{10}.

### 11. Visualization

Visualization of HiC-plot was performed with cooler [https://github.com/open2c/cooler], with each point on the plot representing 80kbp. Circos plot was built with pyCirclize [https://github.com/moshi4/pyCirclize]. The phylogenetic trees were visualized with iTol (Letunic and Bork, 2021). The synteny plots were built with plotsr (Correction to: plotsr: visualizing structural similarities and rearrangements between multiple genomes, 2022), dotplot of genome to genome alignment with dgenies (Cabanettes and Klopp, 2018).

## Results

### 1. Chromosome count

Chromosome number was determined from young anthers of ten plants cultivated from seeds, confirmed by microsatellite genotyping as *B. falcata*. All plants were confirmed to be diploid, with a chromosome count of 2*n* = 14 (Figure 2).

**Figure 2.**
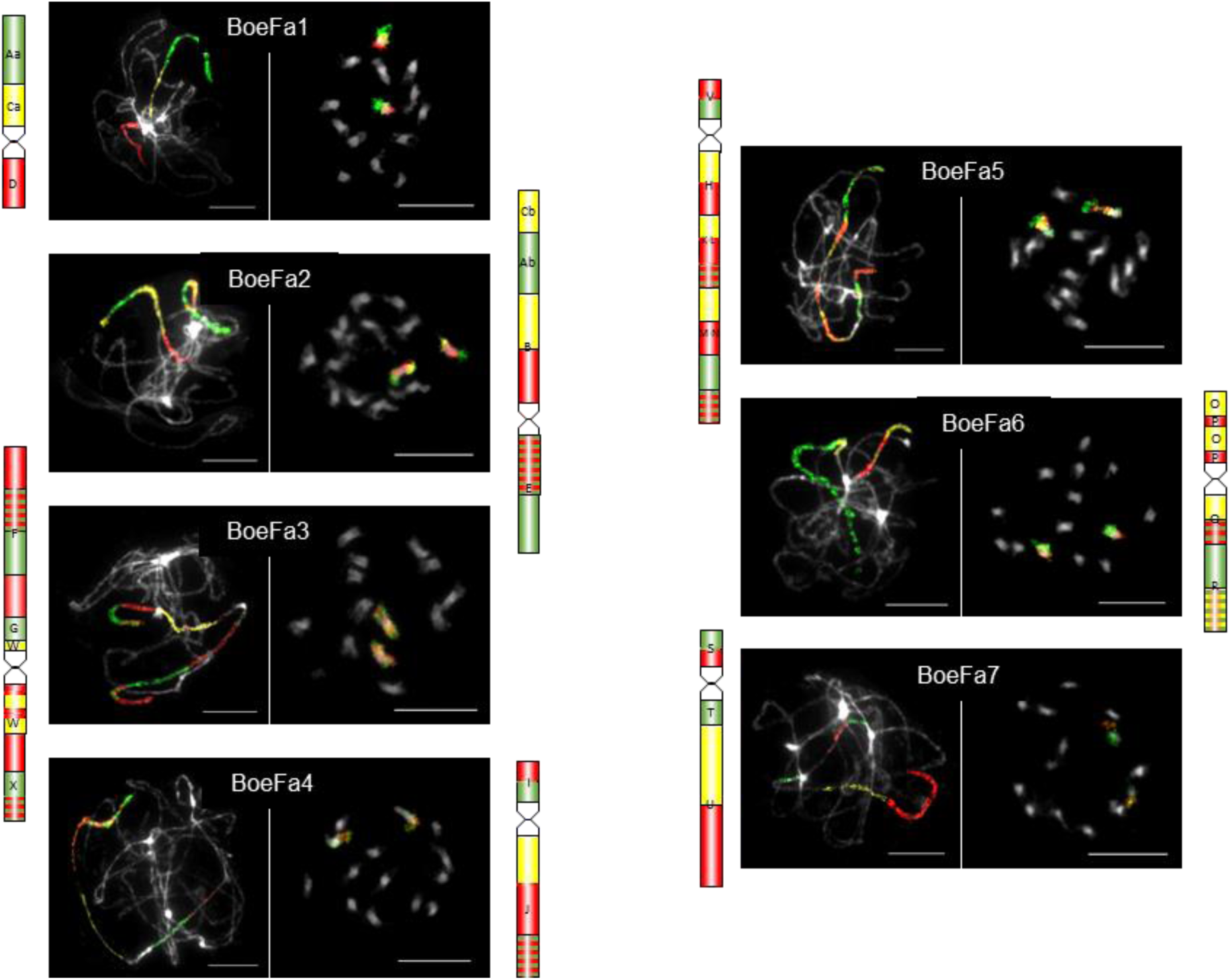
Karyotype structure of *B. falcata* inferred from comparative chromosome painting (CCP). The seven chromosomes (BoeFa1–BoeFa7) were visualized through CCP using *Arabidopsis* BAC contigs as painting probes applied to pachytene, mitotic, and diakinesis chromosome spreads. Painting signals are shown in experimental colors, reflecting the fluorochrome(s) used to label specific genomic blocks (GBs). All chromosomes were counterstained with DAPI and presented as grayscale images. Scale bars, 10 µm.

### 2. Karyotype structure

Bacterial artificial chromosome-based comparative chromosome painting was employed to investigate the genome structure of *B. falcata*. Probes were arranged according to the ancestral Boechereae genome structure (Mandáková et al., 2020), allowing the mapping of all 22 crucifer genomic blocks (Mandáková et al., 2019) onto *B. falcata* chromosomes (BoeFa1–BoeFa7) (Figure 2; 3). Notably, five of the seven *B. falcata* chromosomes (BoeFa1, BoeFa2, BoeFa4, BoeFa5, BoeFa7) exhibit collinearity with their corresponding chromosomes in the ancestral Boechereae genome (Mandáková et al., 2020). In contrast, BoeFa3 contains a pericentric inversion involving genomic block W, while BoeFa6 undergoes a paracentric inversion within its short arm, with breakpoints located within genomic blocks O and P (Figure 2; 3).

**Figure 3.**
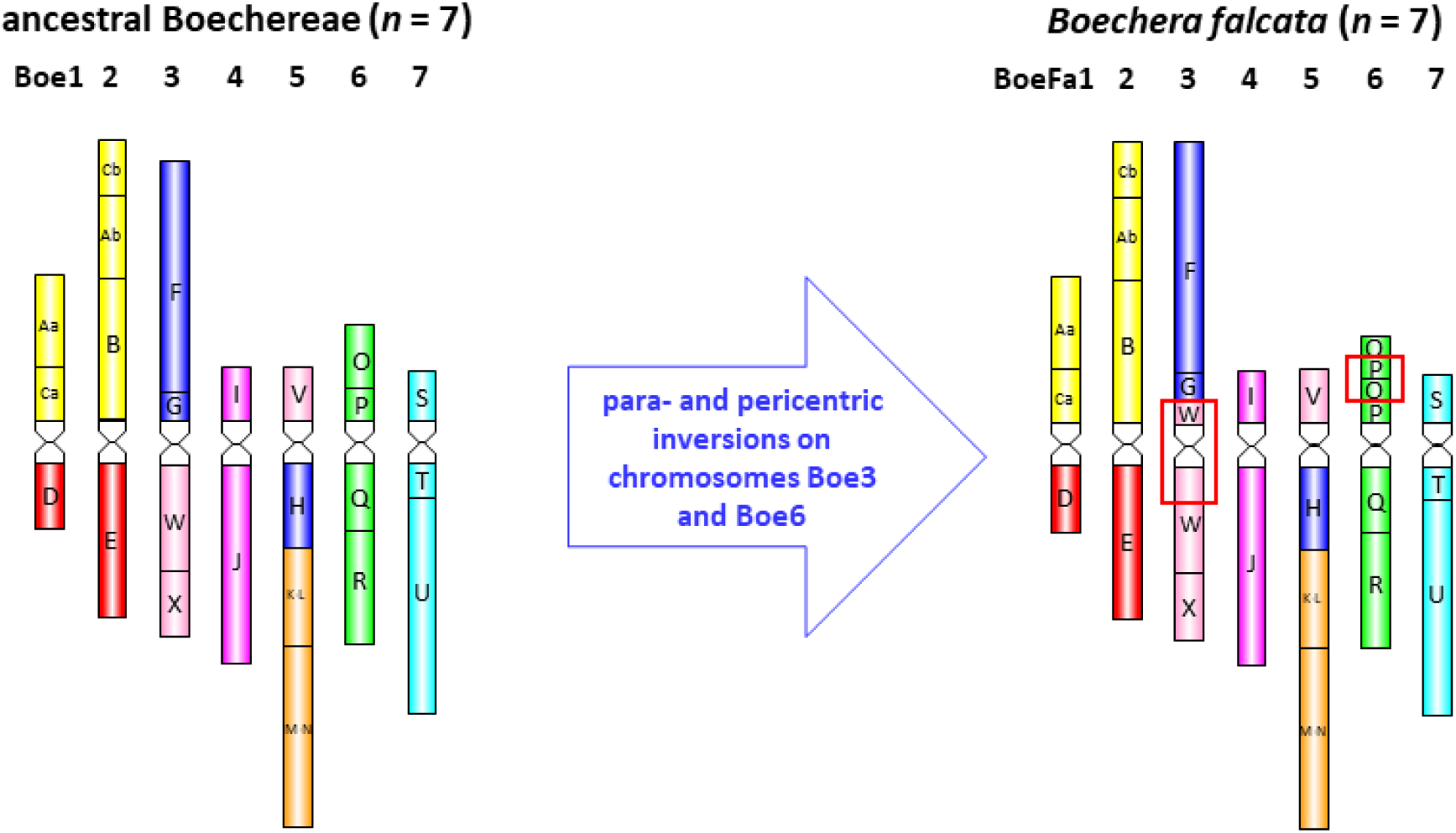
Chromosome evolution of *B. falcata* from the ancestral Boechereae genome (Mandáková et al., 2020).

### 3. k-mer Based Statistics Indicates a High Level of B. falcata Genome Homozygosity

*k*-mer spectrum built by GenomeScope2 software (Ranallo-Benavidez et al., 2020) for both Illumina and HiFi reads is shown in Figure 4. The 23-mer distribution has one major peak at 249× coverage corresponding to diploid 23-mers (shared between homologous chromosomes), also small heterozygous peaks (122× and 64×) were detected (Figure 4). According to the assessment, the genome was highly homozygous with 99,9% of AA k-mers for both Illumina and HiFi reads. The genome size of *B. falcata* was estimated to be close to 185 Mbp by Illumina k-mers and 219 Mbp by HiFi k-mers, which is in the range of the previous estimations of a minimal genome size of ∼200 Mbp in *Boechera* (Anderson et al., 2011). Homozygosity of genomes in *Boechera* is characteristic of sexual reproduction, since all apomicts studied are usually heterozygous, allodiploids or polyploids (Brukhin et al., 2019). Sexuality in *B. falcata* was also confirmed by cytoembryological studies (Vinogradova et al., 2023).

**Figure 4.**
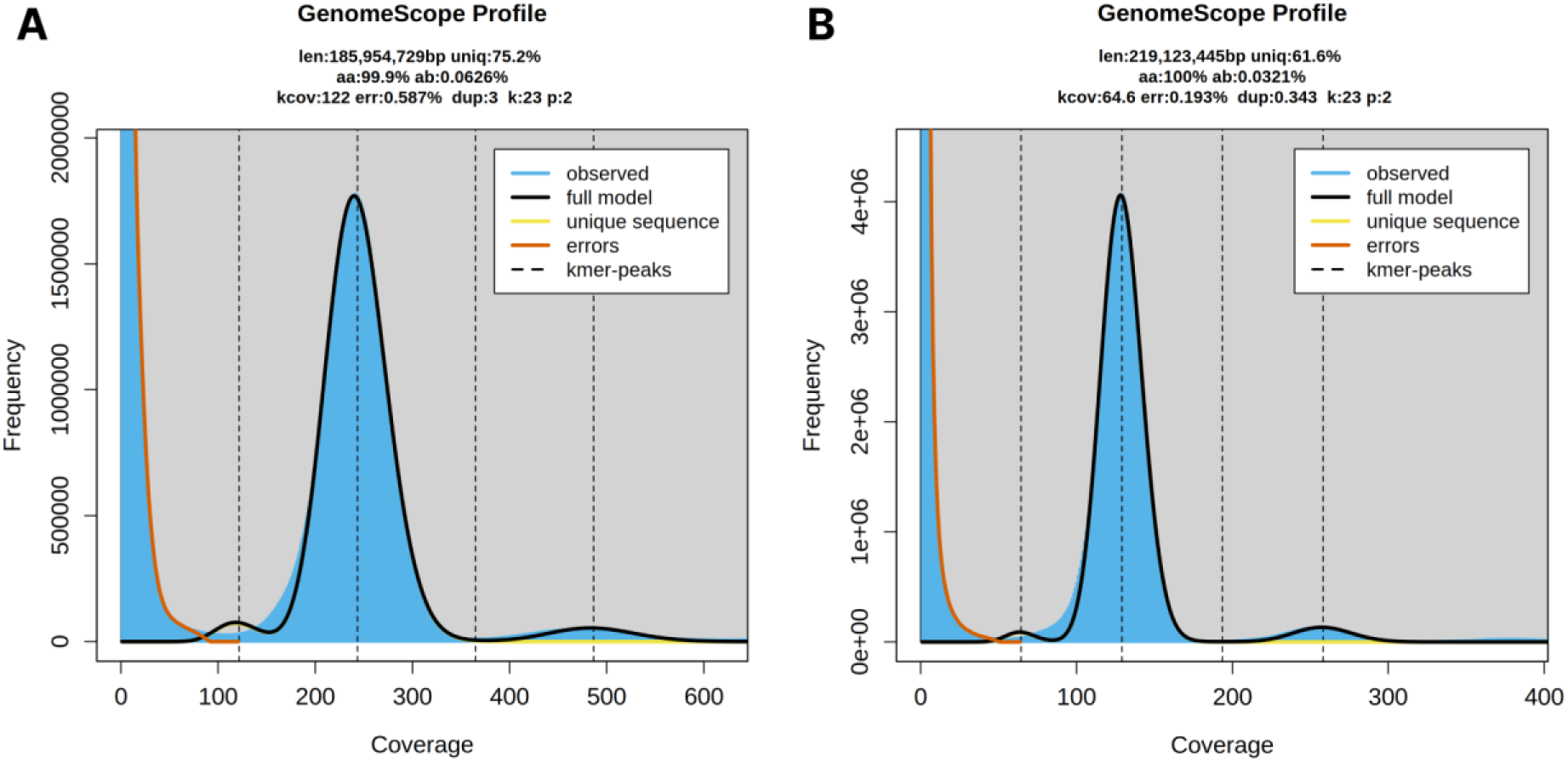
Distribution of 23-mers for detection genome size and evaluation of homo/heterozygosity (A) for Illumina PE reads and (B) HiFi libraries. Only one major peak is present in both Illumina (at 249x coverage) and HiFi (129x) libraries.

### 4. Assessment of the Quality of Genome Assembly Performed by Different Assemblers

To identify the most efficient and correct *Boechera falcata* genome construction, we compared four different assemblers: NextDenovo, flye, hifiasm, and verkko (Table 1). The assemblies varied considerably in their contiguity and quality metrics. NextDenovo assembler demonstrated the highest contiguity with only 50 contigs achieving N50 of 12,407,843 bp, L50 of 7. Evaluation of the assembly completeness was performed using BUSCO, complete BUSCO score is 98.8% (95.8% single-copy, 3.0% duplicated). This assembly also exhibited the highest Merqury QV (62.83) and Merfin QV* (32.99) scores, indicating greater accuracy compared to the other assemblies.

The hifiasm assembler produced the most contigs (1232) and achieved the highest N50 value of 30,472,172 bp. However, it showed a higher number of overrepresented k-mers and lower QV scores compared to the NextDenovo and flye assemblers.

Indeed, all assemblies showed complete BUSCO genes (≥98.7%), indicating high assembly completeness under different assembly strategies. The GC content was similar in all assemblies and ranged from 35.95% to 37.85%. However, based on the obtained metrics, we chose the NextDenovo assembler for further haplotype phasing, polishing, analysis, and improvement of the Hi-C scaffolding due to its favorable balance of continuity, completeness, and accuracy.

### 5. Genome Polishing

After splitting haplotypes to dual assembly with hapdup, both primary and alternate haplotype underwent three rounds of polishing using Illumina reads (Table 2). Each polishing step progressively improved the assembly quality, as evidenced by the decreasing number of missed k-mers and increasing Merqury QV scores. After the final polishing step, the primary haplotype achieved a Merqury QV of 68.04, while the alternative haplotype reached 67.6, indicating high accuracy in both haplotypes.

**Table 2.**
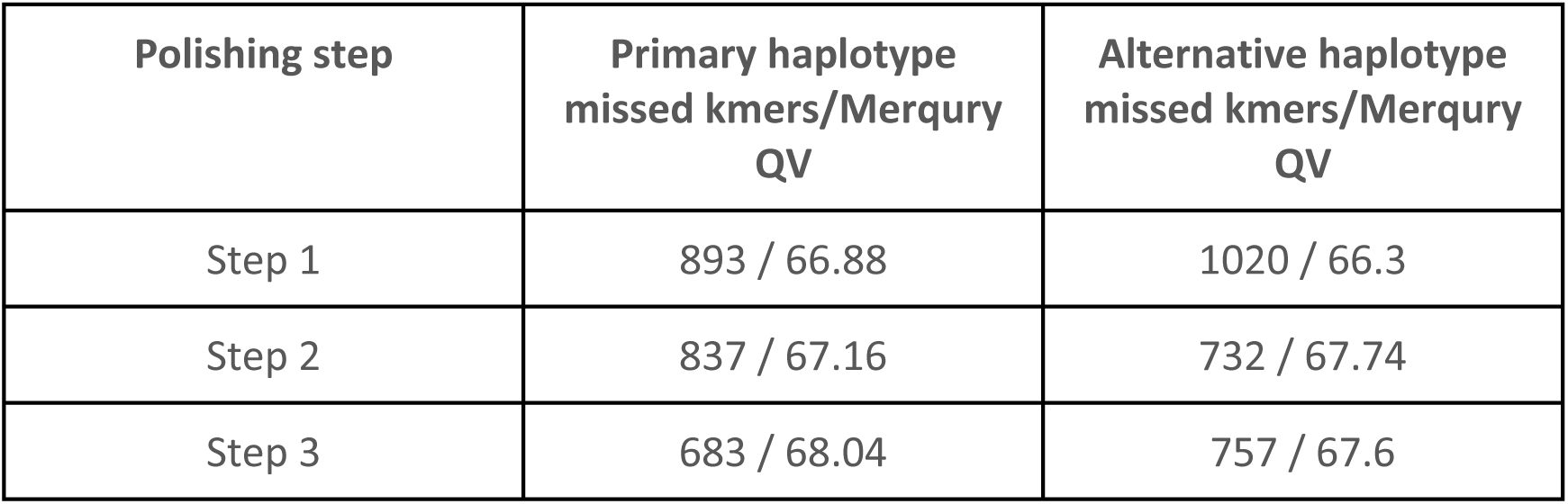
Genome polishing information.

### 6. Hi-C Scaffolding and Chromosome-Level Assembly

Following genome assembly, haplotype phasing, and polishing, we performed Hi-C scaffolding to achieve a chromosome-level assembly. This process resulted in the successful construction of 7 chromosomes, consistent with the known karyotype of *Boechera* species (Fig. 5). The final assembly contained a total of 32 gaps. The assemblies of the primary and alternate haplotypes are available on the NCBI, BioProjrcts: PRJNA1077252 and PRJNA1100512, respectively.

**Figure 5.**
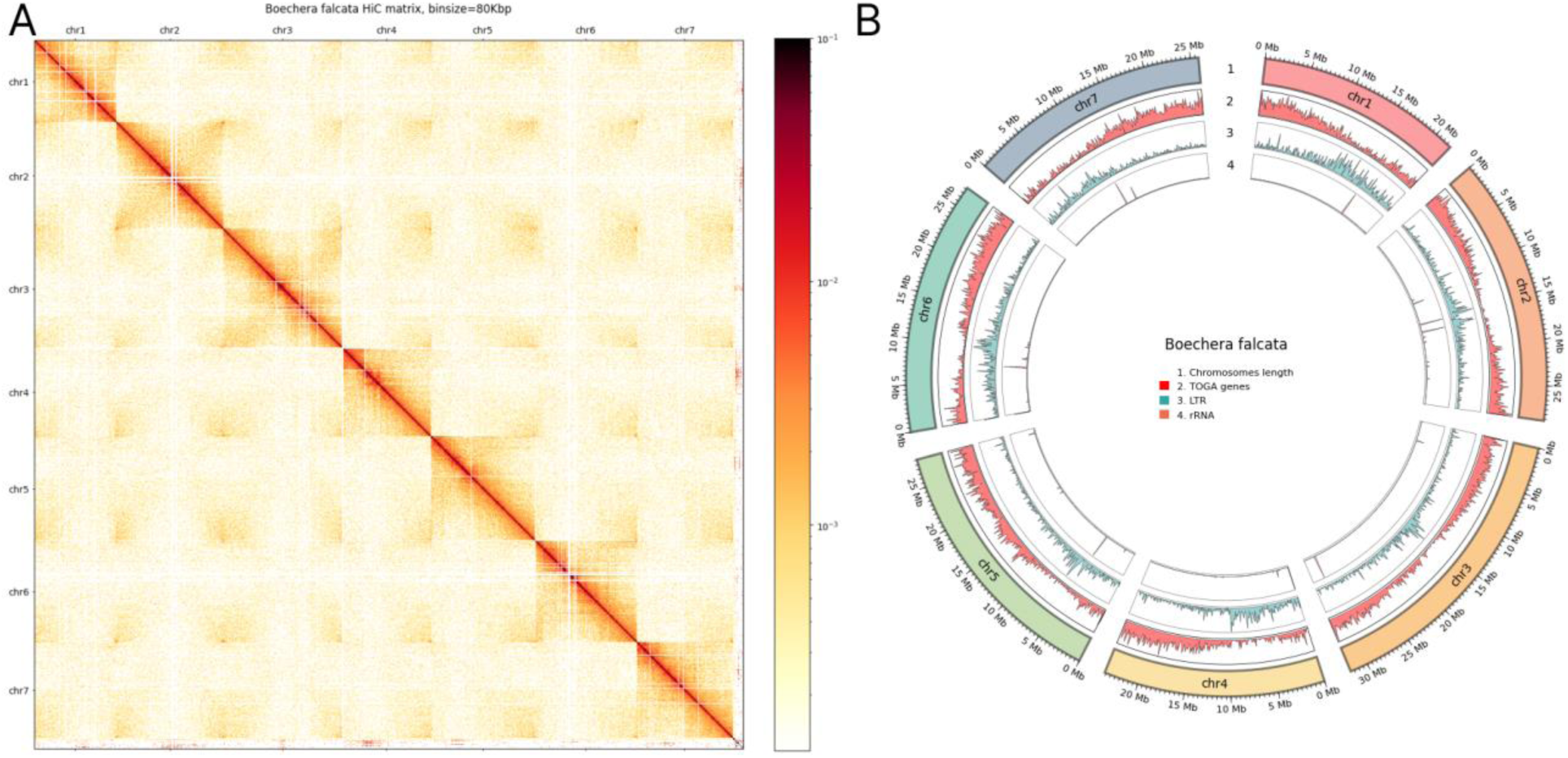
(A) Hi-C contact matrix of *B. falcata* genome at 80Kbp resolution. The seven distinct chromosomes are clearly visible along the diagonal, indicating successful chromosome-level assembly. The intensity of red coloration represents the frequency of interactions between genomic regions, with darker red indicating higher interaction frequencies. (B) Circos plot of the genome assembly. HiC map was build with 80kbp bin size, circos plot includes (1) lengths of chromosomes, (2) frequency of genes in 100kbp bin size, (3) frequency of long tandem repeats in 100kbp bin size and (4) frequency of rRNA genes in the same bin size.

### 7. Analysis of Repetitive Elements

The repeat masking analysis revealed that 38.46% of the genome is made up of repetitive elements (Table 3). Long Terminal Repeat (LTR) retrotransposons were the most common class of repeats, accounting for 19.76% of the genome with 24,870 instances identified. DNA transposons represented the second most prevalent class, masking 6.23% of the genome. Long Interspersed Nuclear Elements (LINEs) and Short Interspersed Nuclear Elements (SINEs) accounted for 2.39% and 0.008% of the genome, respectively. Other repetitive elements, including simple repeats, microsatellites, and RNA genes, comprised for a total of 0.65% of the genome. Additionally, 8.82% of the masked regions were classified as uncharacterized repetitive elements.

**Table 3.** Repeats revealed by RepeatMasker.

| Class of repetitive element | Total length | n of elements | % of genome |
| --- | --- | --- | --- |
| DNA | 11783246 | 17307 | 6.235 |
| LINE | 4527617 | 8007 | 2.39 |
| LTR | 37373250 | 24870 | 19.76 |
| Other (Simple Repeat, Microsatellite, RNA) | 1221953 | 2131 | 0.65 |
| Penelope | 24530 | 259 | 0.01 |
| Rolling Circle | 1123003 | 2785 | 0.59 |
| SINE | 15061 | 234 | 0.01 |
| Unclassified | 16681735 | 47524 | 8.82 |
| **Total** | **72750395** | **103117** | **38.46** |

### 8. Prediction of Protein-Coding Genes

Homology-based prediction yielded a total of 27,516 protein-coding genes predicted. BUSCO analysis demonstrated completeness of annotation, with 99.2% of the 4,596 BUSCO groups present and complete (Table 4).

**Table 4.** Homology-based gene prediction in *Boechera falcata* genome.

|  |  |
| --- | --- |
| Genes | 27516 |
| Exons | 161159 |
| Introns | 130644 |
| mRNAs | 30516 |
| BUSCOs | C:99.2% [S:78.83%, D:20.37%] F:0.13%,<br>M:0.67%, N:4596 |

### 9. Comparative analysis of Boechera falcata and Boechera stricta genomes

Comparative analysis of the *Boechera falcata* and *Boechera stricta* genomes revealed similarities in overall structure, as well as notable differences in assembly quality (Table 5). Both species possess 7 chromosomes and demonstrated highly comparable genome sizes of 189.36 Mb and 190.54 Mb and exhibited similar assembly contiguity metrics N50 of 27.20 Mb and 26.95 Mb for *B. falcata* and *B. stricta* correspondingly. BUSCO genome completeness scores are nearly identical for both species 98.8% and 98.7% for *B. falcata* and *B. stricta* accordingly, as well as gene counts, 27,516 in *B. falcata*, 27,538 in *B. stricta*. However, *B. falcata* demonstrated higher assembly accuracy with higher Merqury QV (68.04 vs. 27.49) and Merfin QV (33.08 vs. 23.32) scores. Both genomes were assembled to chromosome level as diploid assemblies. Thus, combination of high structural similarity and improved accuracy of the *B. falcata* genome assembly provides the basis for a detailed comparative analysis of its genome with other species of the genus *Boechera*.

**Table 5.** Comparison of genome characteristics of *B.falcata* and *B.stricta*.

| Metric | *B.falcata* | *B.stricta* |
| --- | --- | --- |
| Genome length (bp) | 189363209 | 190536481 |
| Chromosomes (x) | 7 | 7 |
| N50 (bp) | 27200968 | 26945205 |
| BUSCO genome | C:98.8%[S:95.8%,D:3.0%],F:0.2%,M:1.0%,n:4596 | C:98.7%[S:95.7%,D:3.0%],F:0.3%,M:1.0%,n:4596 |
| Merquy QV | 68.04 | 27.49 |
| Merfin QV* | 33.08 | 23.32 |
| Genes | 27516 | 27538 |
| BUSCO annotation | C:99.2% [S:78.83%, D:20.37%]<br>F:0.13%, M:0.67%, N:4596 | C:98.98, [S:79.29%,<br>D:19.69%] F:0.13%,<br>M:0.89%, N:4596 |

Analysis of structural variations between the *Boechera falcata* and *Boechera stricta* genomes revealed significant genomic rearrangements (Figure 6). The comparison identified 294 syntenic regions, covering the majority of both genomes (Table 6). A total of 66 inversions were detected, spanning approximately 23.67 Mb in the reference genome (*B. stricta*) and 23.60 Mb in the query genome (*B. falcata*). These inversions represent substantial chromosomal rearrangements among the two species. Large regions of both genomes were found to be non-syntenic: 19.24 Mb are unique to *B. stricta* and 18.36 Mb are unique to *B. falcata*, indicating species-specific genomic content. Figure 6 shows these structural variations on synteny plot (A) and dotplot (B).

**Figure 6.**
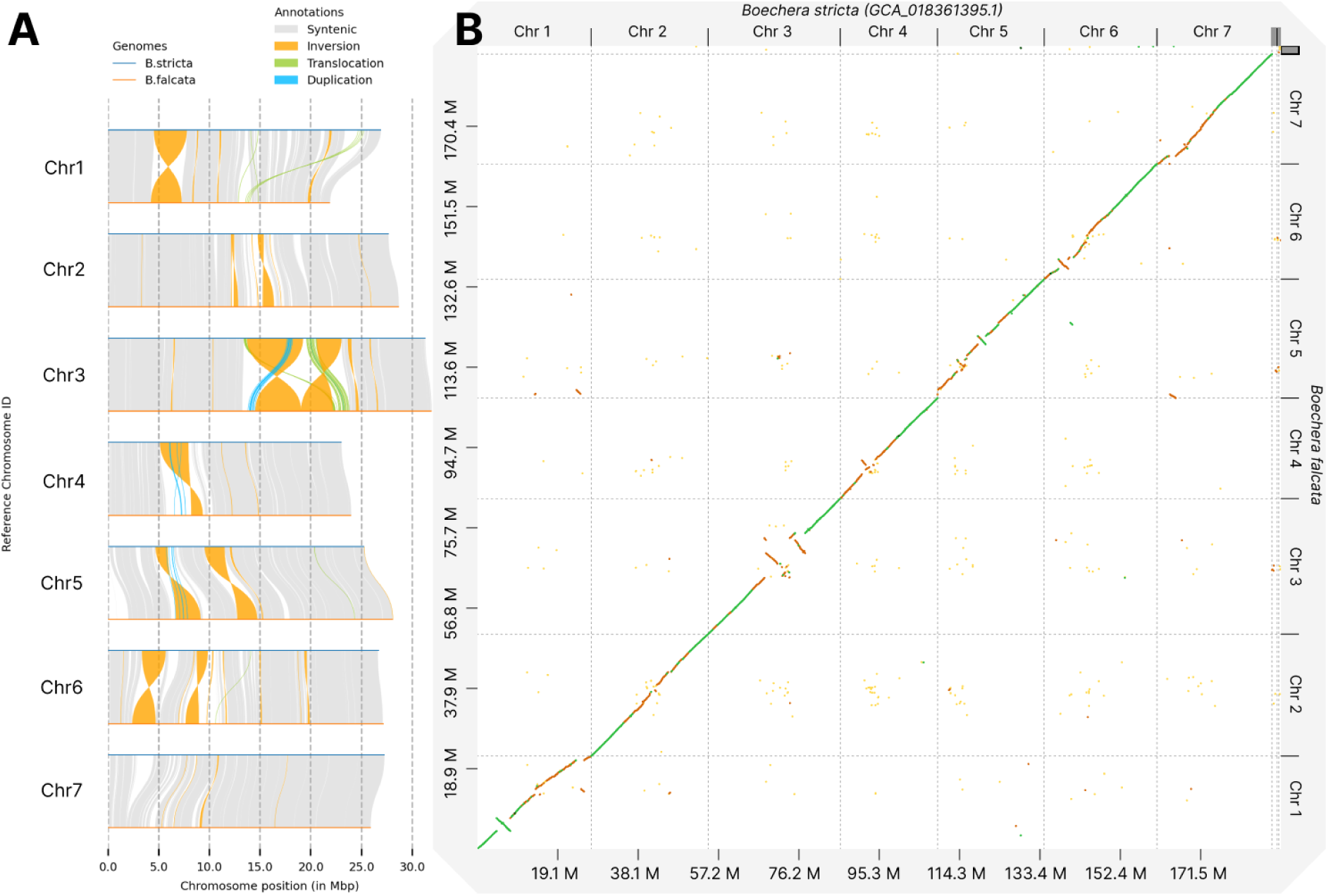
Illustration of the structural variations between *Boechera stricta* and *Boechera falcata* genomes. (A) chromosomal rearrangements, each chromosome of *B. falcata* is aligned to *B. stricta*. The orange regions represent inversions. Gray areas indicate syntenic regions. (B) Scatter plot of genome-wide comparison: chromosomes of *B. stricta* on the x-axis and *B. falcata* on the y-axis. Diagonal green lines represent syntenic regions with high sequence identity (>70%), and yellow regions indicate areas with lower sequence identity (<70%). Prominent large-scale inversions are visible, especially in chromosomes 1, 3, and 5 in (A) and corresponding off-diagonal patterns in (B). The dot plot highlights the overall conservation of chromosomal structure between the two species and also highlights significant inversions and areas of differing sequence identity across the genomes.

**Table 6.** Synteny analysis characteristics.

| Variation_type | Count | Length_ref (*B.stricta*) | Length_qry (*B.falcata*) |
| --- | --- | --- | --- |
| Syntenic regions | 294 | 141724185 | 141383384 |
| Inversions | 66 | 23669784 | 23602144 |
| Translocations | 348 | 3730041 | 3629426 |
| Duplications (*B.stricta*) | 150 | 476406 | - |
| Duplications (*B.falcata*) | 359 | - | 1366155 |
| Not aligned (*B.stricta*) | 801 | 19235167 | - |
| Not aligned (*B.falcata*) | 1030 | - | 18357653 |

### 10. B. falcta Falls into the Homozygous Sex Allele Clade on the APOLLO Phylogenetic Tree

The *APOLLO* gene, encoding the NEN3 exonuclease, is an important apomixis- associated gene in *Boechera* (Corral et al., 2013; Brukhin et al., 2019). The *APOLLO* locus has previously been shown to have multiple alleles (Corral et al., 2013; Kliver et al., 2018). All studied apomictic plants carry at least one of the “apo” alleles, and the sexual accessions are homozygous for the “sex” alleles.

We decided to further explore the *APOLLO* locus in the *B. falcata* genome. The phylogenetic analysis of the *APOLLO* gene revealed a clear division between two groups of *Boechera* species, in accordance with their mode of reproduction (Figure 7), as was previously described (Kliver et al. 2018). The tree shows two distinct clades within the *Boechera* genus, with strong bootstrap support (100) for the split between these groups. One clade comprises apomictic *Boechera spp.,* including apomictic *Boechera* hybrid M4B, whose genome we assembled to chromosome scale previously (NCBI BioProject PRJNA774175), which forms a well-supported monophyletic group. The other clade comprises *B. falcata*, *B. arcuata*, *B. retrofracta*, *B. stricta*, and several *Boechera spp.*, along with multiple *B. spatifolia* accessions, which reproduce sexually. Notably, *B. falcata* is positioned within this second clade, clustering closely with sexual *B. arcuata* and *Boechera spp.* (KF705572.1). This study demonstrates clear phylogenetic differentiation of the *APOLLO* gene between the two groups of *Boechera* species, with the *B. falcata* group included in the clade that along with *B. stricta* and other known sexual species.

**Figure 7.**
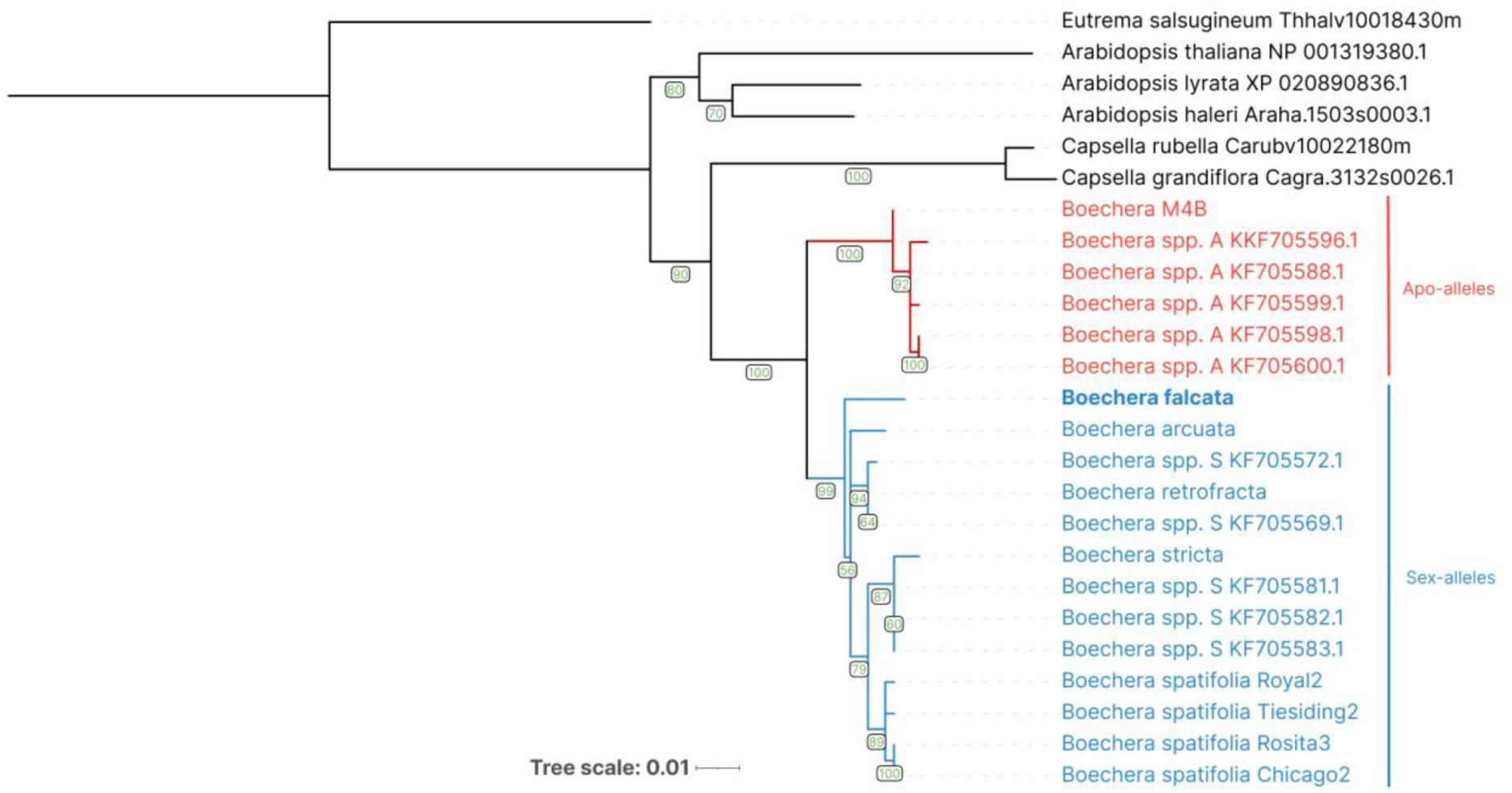
Phylogenetic tree of *APOLLO* gene alleles in *Boechera* and related species. Sequences of *Eutrema salsugineum, Arabidopsis thaliana, A. lyrata, A. haleri, Capsella rubella,* and *C. grandiflora* were used as outgroups. The two main clades within *Boechera* are highlighted: apomictic species in red and sexual species in blue including *B. falcata*. The numbers in the nodes represent bootstrap support values.

### 11. Chloroplast and mitochondrial genome assembly and annotation

To facilitate further evolutionary relationship study of non-model *Boechera* species, phylogenetic researches, and species identifications we performed *B. falcata* chloroplast and mitochondrial genomes assembly and annotation (Figure 8; 9). 79 unique CDS, 30 tRNAs and 4 rRNA were identified in the chloroplast genome. Large single-copy (LSC) part and small single-copy (SSC) part of the chloroplast DNA occupy 84428 bp and 17952 bp respectively. GC contents is 36.41%. All these indicators are consistent with those in the chloroplast genomes of other *Boechera* species as shown in Table 7.

**Figure 8.**
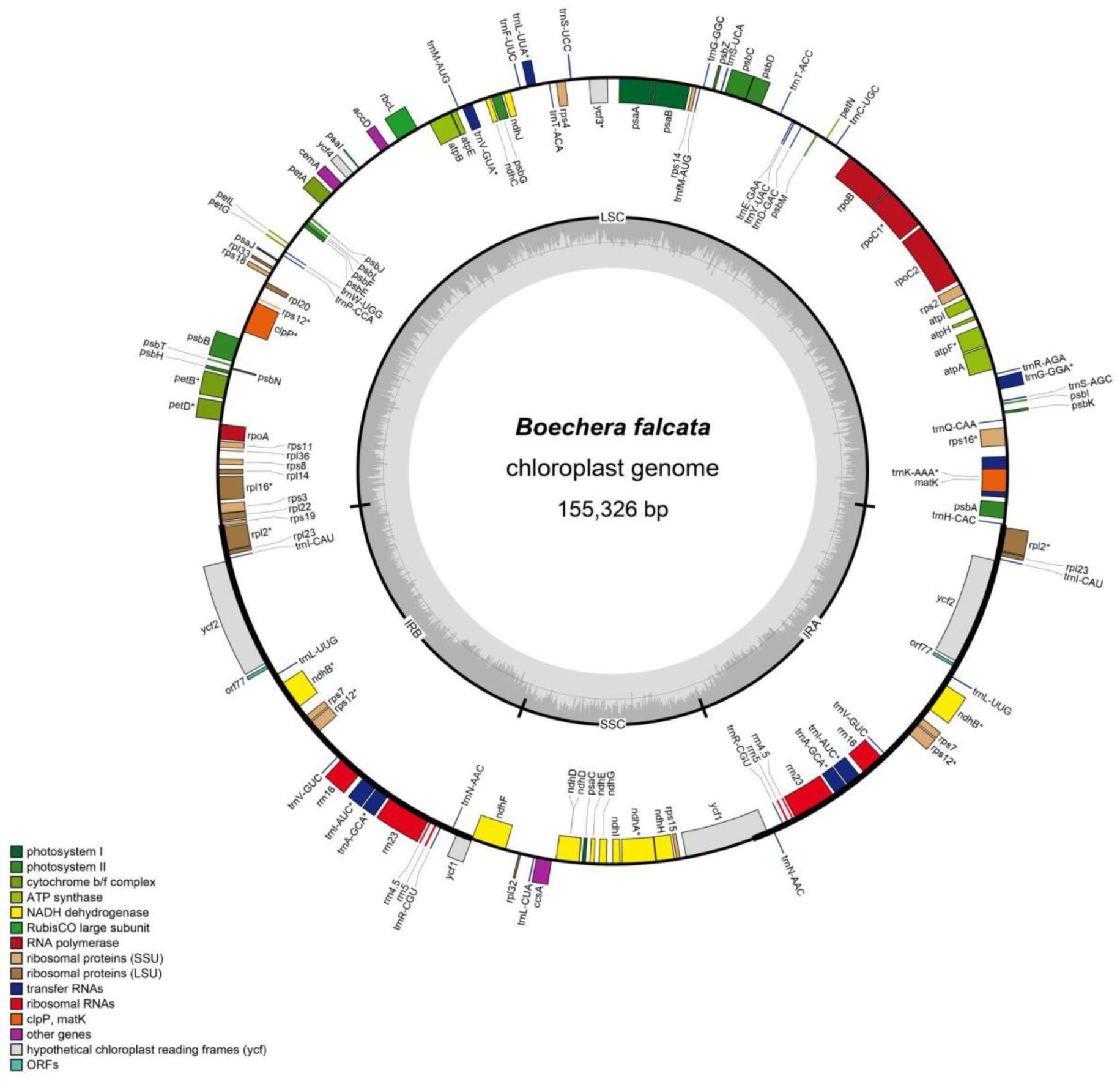
Circular representation of *B. falcata* chloroplast genome. Gray circle inside is GC% content.

**Table 7.** Comparison of several *Boechera* species chloroplast genomes.

| Species name | RefSeq accession number | Genome size (bp) | LSC Length (bp) | SSC Length (bp) | IR Length (bp) | Number of genes |  |  | GC(%) |  |  |  |
| --- | --- | --- | --- | --- | --- | --- | --- | --- | --- | --- | --- | --- |
|  |  |  |  |  |  | CDS | tRNAs | rRNAs | Total genome | LSC | SSC | IR |
| *Boechera falcata* | chloroplast | 155326 | 84428 | 17952 | 26473 | 79 | 30 | 4 | 36.41 | 34.20 | 29.42 | 42.33 |
| *Boechera suffrutescens* | NC_049600.1 | 155114 | 84285 | 17915 | 26457 | 79 | 30 | 4 | 36.43 | 34.18 | 29.52 | 42.36 |
| *Boechera stricta* | NC_049599.1 | 155033 | 84237 | 17884 | 26456 | 79 | 30 | 4 | 36.39 | 34.15 | 29.38 | 42.33 |
| *Boechera pulchra* | NC_049598.1 | 155348 | 84483 | 17951 | 26457 | 79 | 30 | 4 | 36.42 | 34.17 | 29.49 | 42.37 |
| *Boechera puberula* | NC_049597.1 | 155011 | 84189 | 17928 | 26447 | 79 | 30 | 4 | 36.48 | 34.26 | 29.63 | 42.35 |
| *Boechera laevigata* | NC_049596.1 | 155083 | 84266 | 17891 | 26463 | 79 | 30 | 4 | 36.45 | 34.22 | 29.48 | 42.33 |
| *Boechera davidsonii* | NC_049594.1 | 154935 | 84107 | 17914 | 26457 | 79 | 30 | 4 | 36.46 | 34.24 | 29.57 | 42.33 |
| *Boechera breweri* | NC_049593.1 | 155147 | 84347 | 17886 | 26457 | 79 | 30 | 4 | 36.44 | 34.21 | 29.56 | 42.33 |

Length of the mitochondrial DNA in *B. falcata* is 292 083 bp and GC content is 44.90% (Figure 9). Comparative data on the mitochondrial genome size and GC content in *B. falcata, B. stricta*, and *A. thaliana* are shown in table 8. Table 9 indicates the type and functions of the genes identified in mitochondrial DNA of *B. falcata, B. stricta*, and *A. thaliana.* All in all 33 protein-coding genes, 17 tRNAs and 3 rRNA were annotated in the mitochondrial genome of *B. falcata*.

**Figure 9.**
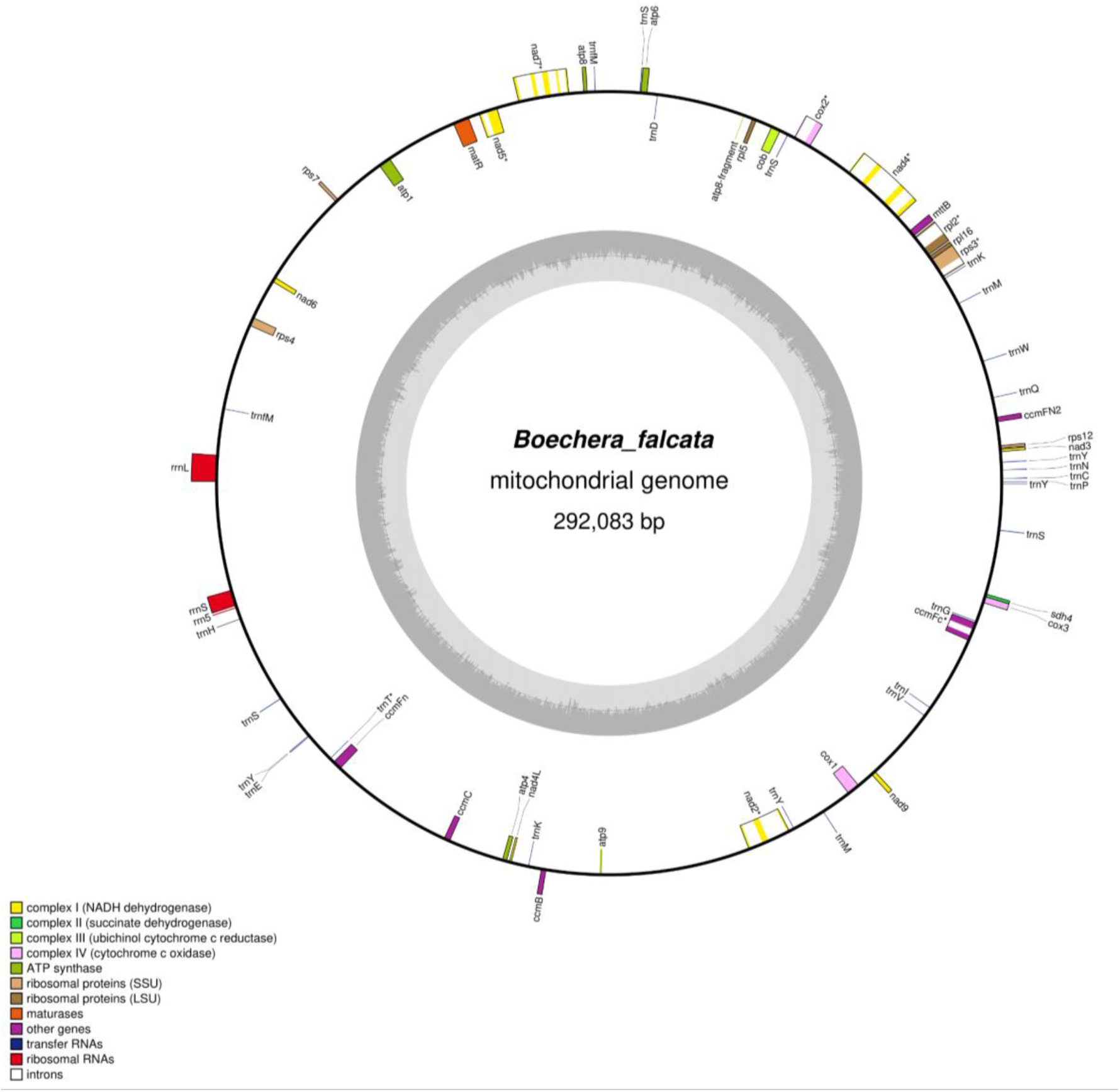
Circular diagram of *B. falcata* mitochondrial genome. Gray circle inside shows % of GC content.

**Table 8.** Comparative data on mitochondrial genome size and GC content in *B. falcata, B. stricta*, and *A. thaliana*.

| Species name | RefSeq accession number | Genome size (bp) | GC content % |
| --- | --- | --- | --- |
| *Boechera falcata* | mithochondrion | 292083 | 44.90 |
| *Boechera stricta* | NC_042143.1 | 271601 | 44.93 |
| *Arabidopsis thaliana* | NC_037304.1 | 367808 | 44.79 |

**Table 9.** Characteristics of the genes identified in mitochondrial DNA of *B. falcata, B. stricta*, and *A. thaliana*.

| Group of genes | Name of genes |  |  |
| --- | --- | --- | --- |
|  | *Boechera falcata* | *Boechera stricta* | *Arabidopsis thaliana* |
| ATP synthase | atp1, atp4, atp6, atp8, atp9 | atp1, atp4, atp6, atp8, atp9 | atp1, atp4, atp6, atp8, atp9 |
| Cytochrome c biogenesis | ccmB, ccmC, ccmFc, ccmFn, ccmFN2 | ccmB, ccmC, ccmFc, ccmFn | ccmB, ccmC, ccmFC, ccmFN1, ccmFN2 |
| Ubiquinol cytochrome c reductase | cob | cob | cob |
| Cytochrome c oxidase | cox1, cox2, cox3 | cox1, cox2, cox3 | cox1, cox2, cox3 |
| Maturases | matR | matR | matR |
| Transport membrane protein | mttB | mttB | mttB |
| NADH dehydrogenase | nad1, nad2, nad3, nad4, nad4L, nad5, nad6, nad7, nad9 | nad1, nad2, nad3, nad4, nad4L, nad5, nad6, nad7, nad9 | nad1, nad2, nad3, nad4, nad4L, nad5, nad6, nad7, nad9 |
| Large subunit of ribosome | rpl2, rpl5, rpl16 | rpl2, rpl5, rpl16 | rpl2, rpl5, rpl16 |
| Small subunit of ribosome | rps3, rps4, rps7, rps12 | rps3, rps4, rps12 | rps3, rps4, rps7, rps12 |
| Succinate dehydrogenase | sdh4 | sdh4 | - |
| Ribosomal RNAs | rrn5, rrn18, rrn26 | rrn5, rrn18, rrn26 | rrn5, rrn18, rrn26 |
| Transfer RNAs | trnC, trnD, trnE, trnG, trnH, trnI, trnK, trnM, trnN, trnP, trnQ, trnS, trnT, trnV, trnW, trnY, trnfM | trnC, trnD, trnE, trnG, trnH, trnI, trnK, trnM, trnN, trnP, trnQ, trnS, trnW, trnY, trnfM | trnC, trnD, trnE, trnG, trnH, trnI, trnK, trnM, trnN, trnP, trnQ, trnS, trnW, trnY, trnfM |

### 12. Phylogeny of B. falcata and Its Relationship with Other Boechera and Brassicaceae species

A phylogenetic tree of Brassicacae constructed based on chloroplast DNA (Figure 10) shows that *B. falcata* is nested among the North American *Boechera* species, with the closest relatives of *B. falcata* being *B. sufrutescens* and *B. pulchra*.

**Figure 10.**
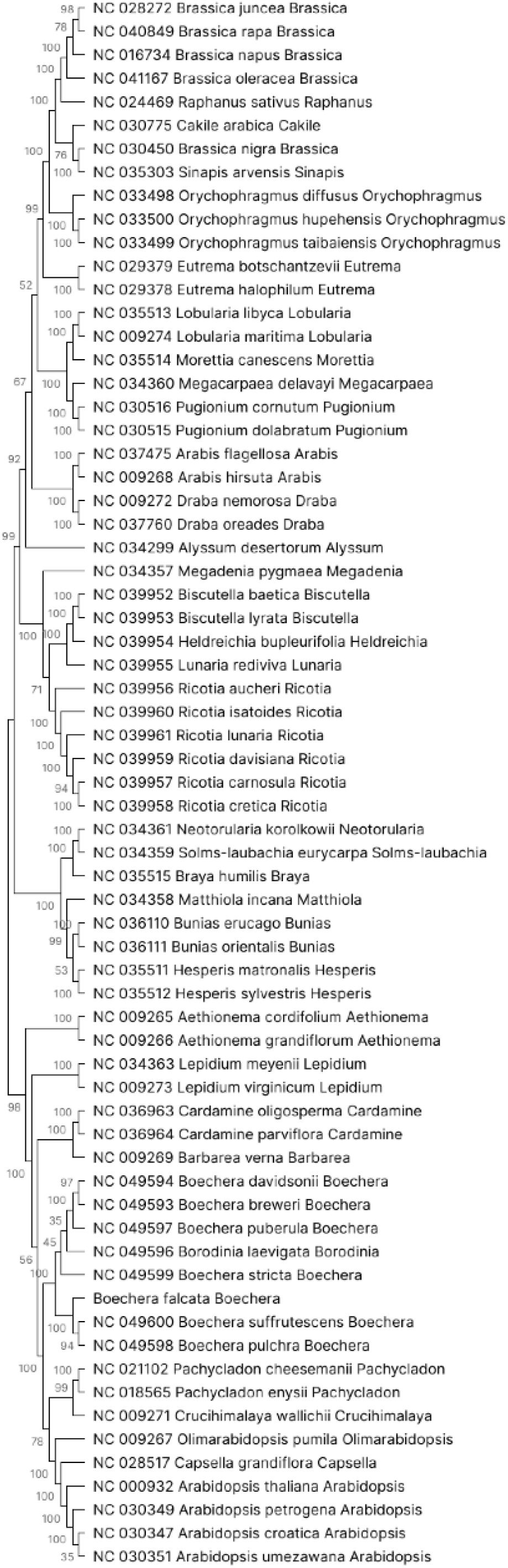
Phylogenetic tree of Brassicaceae species based on the chloroplast genes.

The combination of Angiosperms353 and Brassicaceae764 probe sets revealed a comprehensive evolutionary relationship within the Boechereae tribe (Figure 11). The tree strongly supports the deep divergence between *Boechera* sensu stricto and the non- *Boechera* Boechereae (NBB) clade, consistent with previous studies (Hay et al., 2023). Within *Boechera* s.s., we observed three main core groups (Core *Boechera* I, II, and III). *Boechera falcata*, the focus of our study, is positioned within the Core *Boechera* III, sister to what Hay et al. (2023) called the Boreal clade. This placement provides new insights into the phylogenetic relationships of *B. falcata* within the broader *Boechera* genus. The tree topology and strong bootstrap support of many nodes demonstrate the power of combining these probe sets for resolving relationships within this complex group.

**Figure 11.**
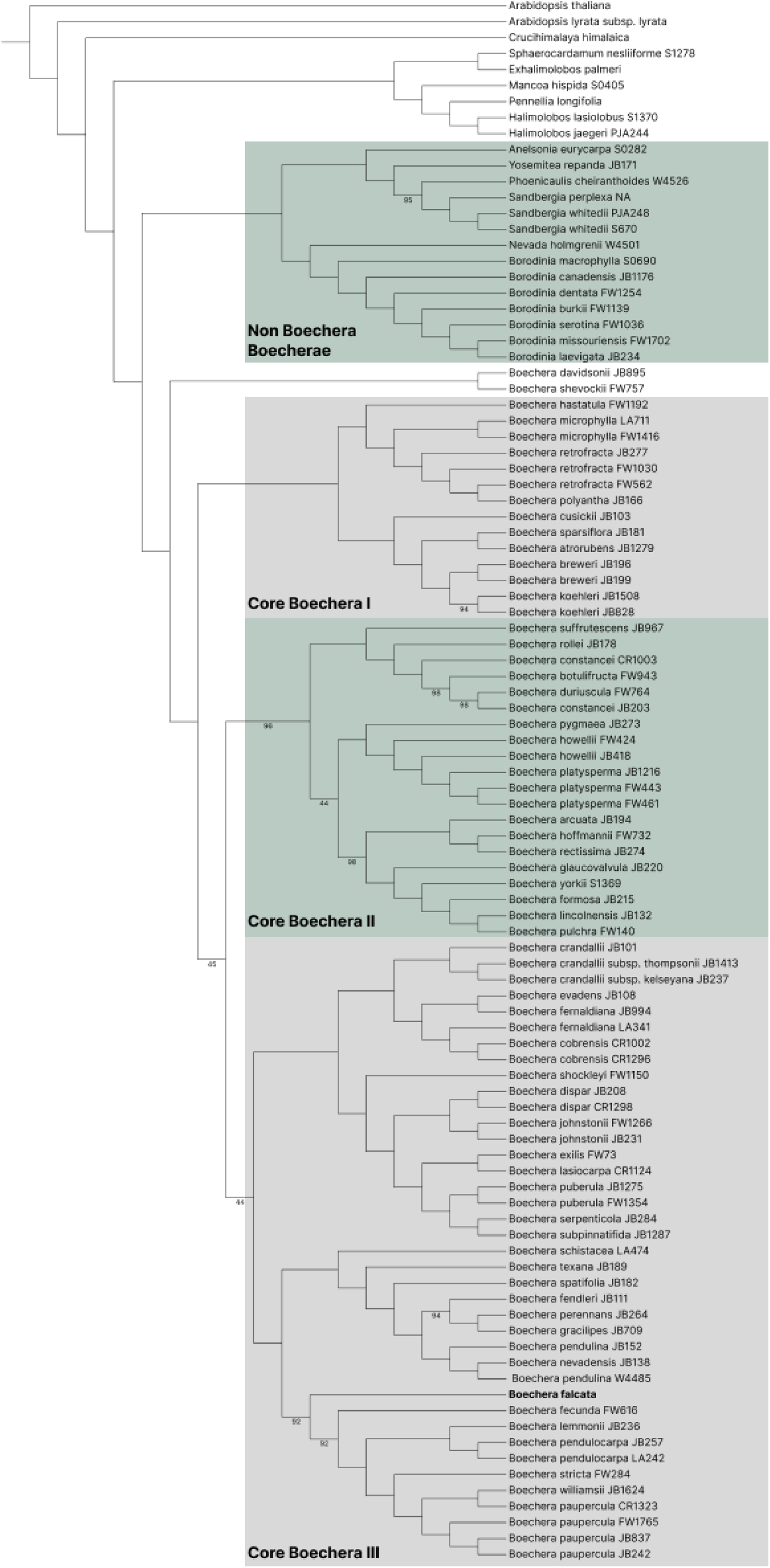
Phylogenetic tree of Boechereae and related species based on combined Angiosperms353 and Brassicaceae764 probe sets. The tree is rooted with outgroup species including *Arabidopsis thaliana* and *Schrenkiella parvula*. Major clades are highlighted. *Boechera falcata* is indicated in bold. Numbers at nodes represent bootstrap support values <100.

## Conclusion

To conclude, we present a high-quality genome assembly of the only *Boechera* species native to Eurasia, (the Far Eastern *B. falcata*) to the chromosome level, as well as an assembly of chloroplast and mitochondrial DNA. The highly homozygous genome, the presence of only sex alleles of the *APOLLO* gene and previously obtained cytoembryological data suggest that this species reproduces only sexually. It likely came to the Eurasian continent from North America across the Bering Land Bridge during the Pleistocene glacial maxima. In terms of genome morphology, structure and size, as well as chromosome arrangement and organelle DNA organization, this species is quite close to its North American sexual relatives from the Boechereae tribe. Using molecular probes, it was possible to confirm the placement of *B. falcata* within tribe Boechereae, namely in the Core *Boechera* III group. A phylogenetic tree constructed based on the chloroplast DNA confirmed the phylogenetic analysis using molecular probes. Within our limited sampling for the chloroplast DNA phylogeny, *B. falcata* appears most closely related to *B. suffrutescens* and *B. pulchra*. Finally, the availability of the genome of the only Eurasian *Boechera* species will help studies of the evolution and phylogeny of Brassicaceae species, as well as apomixis researchers to unravel the complex hybridization events that form *Boechera* apomicts.

## Acknowledgement

The work was carried out within the framework of the state assignment of the Komarov Botanical Institute RAS No. 124013100862-0 “Polyvariance of morphogenetic programs for the development of plant reproductive structures, regulation of morphological processes in vivo and in vitro” (2024-2028) and RFBR grant No. 20-54-46002 “Phylogenetic and functional analysis of genes associated with apomixis in representatives of the *Boechera* genus” to VB. This work was further supported by the Czech Science Foundation (project 24- 11371S to TM). Seeds and photographs of *B. falcata* plants were kindly provided by Dr. N.V. Sinelnikova from the Institute of Biological Problems of the North, Far Eastern Branch RAS.

